# Fast Evolution of SOS-Independent Multi-Drug Resistance in Bacteria

**DOI:** 10.1101/2022.03.29.486198

**Authors:** Le Zhang, Yunpeng Guan, Yuen Yee Cheng, Nural N. Cokcetin, Amy L. Bottomley, Andrew Robinson, Elizabeth J. Harry, Antoine van Oijen, Qian Peter Su, Dayong Jin

## Abstract

The killing mechanism of many antibiotics involves the induction of DNA damage, either directly or indirectly, which activates the SOS response. RecA, the master regulator of the SOS response, has shown to play a central role in the evolution of resistance to fluoroquinolones, even after short-term exposure. While this paradigm is well established for DNA-damaging antibiotics, it remains unclear whether β-lactams elicit similar resistance dynamics or depend on RecA and SOS-mediated mechanisms. In this study, we observed a rapid and stable evolution of β-lactam resistance (20-fold MIC increase within 8 hours) in *Escherichia coli* lacking RecA after a single exposure to ampicillin. Contrary to expectation, this resistance emerged through an SOS-independent mechanism involving two distinct evolutionary forces: increased mutational supply and antibiotic-driven selection. Specifically, we found that RecA deletion impaired DNA repair and downregulated base excision repair pathways, while concurrently repressing the transcription of antioxidative defence genes. This dual impairment led to excessive accumulation of reactive oxygen species (ROS), which in turn promoted the emergence of resistance-conferring mutations. While ampicillin treatment did not alter survival, it selectively enriched for rare mutants arising in the RecA-deficient and ROS-elevated background. Collectively, our findings demonstrate that this oxidative environment, together with compromised DNA repair capacity, increases genetic instability and creates a selective landscape favouring the expansion of resistant clones. These results highlight the repair-redox axis as a key determinant of bacterial evolvability under antimicrobial stress.

## Introduction

Addressing bacterial infections caused by emerging and drug-resistant pathogens represents a major global health priority. Bactericidal antibiotics can exert their effects on cells by directly or indirectly causing DNA damage or triggering the production of highly destructive hydroxyl radicals (1–3). This, in turn, initiates a protective mechanism known as the SOS response, which enables bacterial survival against the lethal impacts of antibiotics by activating intrinsic pathways for DNA repair (4–7). The activation of DNA repair processes relies on specific genes, such as *recA*, which encodes a recombinase involved in DNA repair, and *lexA*, a repressor of the SOS response that can be inactivated by RecA (8).

Studies have demonstrated that a single exposure to fluoroquinolones, a type of antibiotic that induces DNA breaks and triggers the SOS response, leads to the development of bacterial resistance in *Escherichia coli* (*E. coli*) through a RecA and SOS response-dependent mechanism (9). Given the crucial role of RecA in the SOS response, inhibiting RecA activity to deactivate the SOS response presents an appealing strategy for preventing the evolution of bacterial resistance to antibiotics (10). Similarly, exposure to fluoroquinolones induces the SOS response and mutagenesis in *Pseudomonas aeruginosa*, and the deletion of *recA* in this pathogen results in a significant reduction in resistance to fluoroquinolones (11).

Unlike fluoroquinolones, β-lactam antibiotics induce a RecA-dependent SOS response in *E. coli* through impaired cell wall synthesis, mediated by the DpiBA two-component signal system (12). The development of antibiotic resistance, triggered by exposure to β-lactams, has been extensively investigated using the cyclic adaptive laboratory evolution (ALE) method. Mutations that arise during cyclic ALE experiments are attributed to errors occurring during continued growth, necessitating multiple rounds of β-lactam exposure to drive the evolution of resistance in *E. coli* cells (13,14). However, the precise roles of RecA and SOS responses in the development of resistance under short-term β-lactam antibiotics exposure remain unclear.

Recently, there has been a growing interest in understanding the impact of the stress-induced accumulation of reactive oxygen species (ROS) on bacterial cells (15,16). While exploring methods to harness ROS-mediated killing has the potential to enhance the effectiveness of various antibiotics (17–19), the role of ROS in antimicrobial activity has become a topic of controversy following challenges to the initial observations (20,21). The generation of ROS has been found to contribute to the development of multidrug resistance, as prolonged exposure to antibiotics in cyclic ALE experiments is known to generate ROS, leading to DNA damage and increased mutagenesis (22,23). Nevertheless, there is still limited knowledge regarding the consequences of ROS accumulation in bacteria when the activity of RecA or the SOS response is suppressed.

Here, we report that a single exposure to β-lactam antibiotics can rapidly drive the evolution of multidrug resistance in *E. coli* lacking RecA. This process reflects a two-step evolutionary mechanism: RecA deficiency increases mutational supply by impairing DNA repair, repressing antioxidant gene expression, and promoting ROS accumulation; subsequently, antibiotic pressure selectively enriches resistant variants from this hypermutable population.

## Results

### Single β-lactam exposure accelerates resistance evolution in the *recA* mutant strain through a selection-driven mechanism

To investigate the impact of the SOS response on bacterial evolution towards β-lactam resistance, we generated a *recA* mutant strain (Δ*recA*) from the *E. coli* MG1655 strain. Initially, we conducted an ALE experiment using a slightly modified treatment protocol (24) on the wild type and Δ*recA* strains. During a period of three weeks, the cells were subjected to cycles of ampicillin exposure for 4.5 hours at a concentration of 50 µg/mL (10 times the MIC) each day (Figure 1-figure supplement 1A) (25). As anticipated based on previous studies, the intermittent ampicillin treatments over the course of three weeks resulted in the evolution of antibiotic resistance in the wild type strain (Figure 1-figure supplement 1B). However, a significantly accelerated development of resistance was found in the Δ*recA* strain, with the average time to resistance reduced to 2 days (Figure 1-figure supplement 1B). More importantly, we observed that resistance even emerged in the Δ*recA* strain after a single exposure for 8 hours to ampicillin (Fig. 1A-C). Meanwhile, after 8 hours of treatment with 50 μg/mL ampicillin, the survival rates of both wild type and Δ*recA* strain were consistent (Figure 1-figure supplement 2). To ensure that the emergence of resistance we observed was not illusory due to technical issues during the *recA* knockout process, we employed another Δ*recA* strain (JW2669-1) provided by the Coli Genetic Stock Centre (CGSC) with the same killing procedure. The results from both bacterial strains were consistent (Figure 1-figure supplement 3A and B). To further investigate, we treated both the wild type and Δ*recA* cells with other β-lactams, including penicillin G and carbenicillin, at concentrations equivalent to 10 times the MIC (1 mg/mL and 200 µg/mL, respectively) for 8 hours (26,27). Consistently, these treatments also led to a fast evolution of antibiotic resistance in the Δ*recA* strain (Figure 1-figure supplement 4A and B).

**Figure 1.**
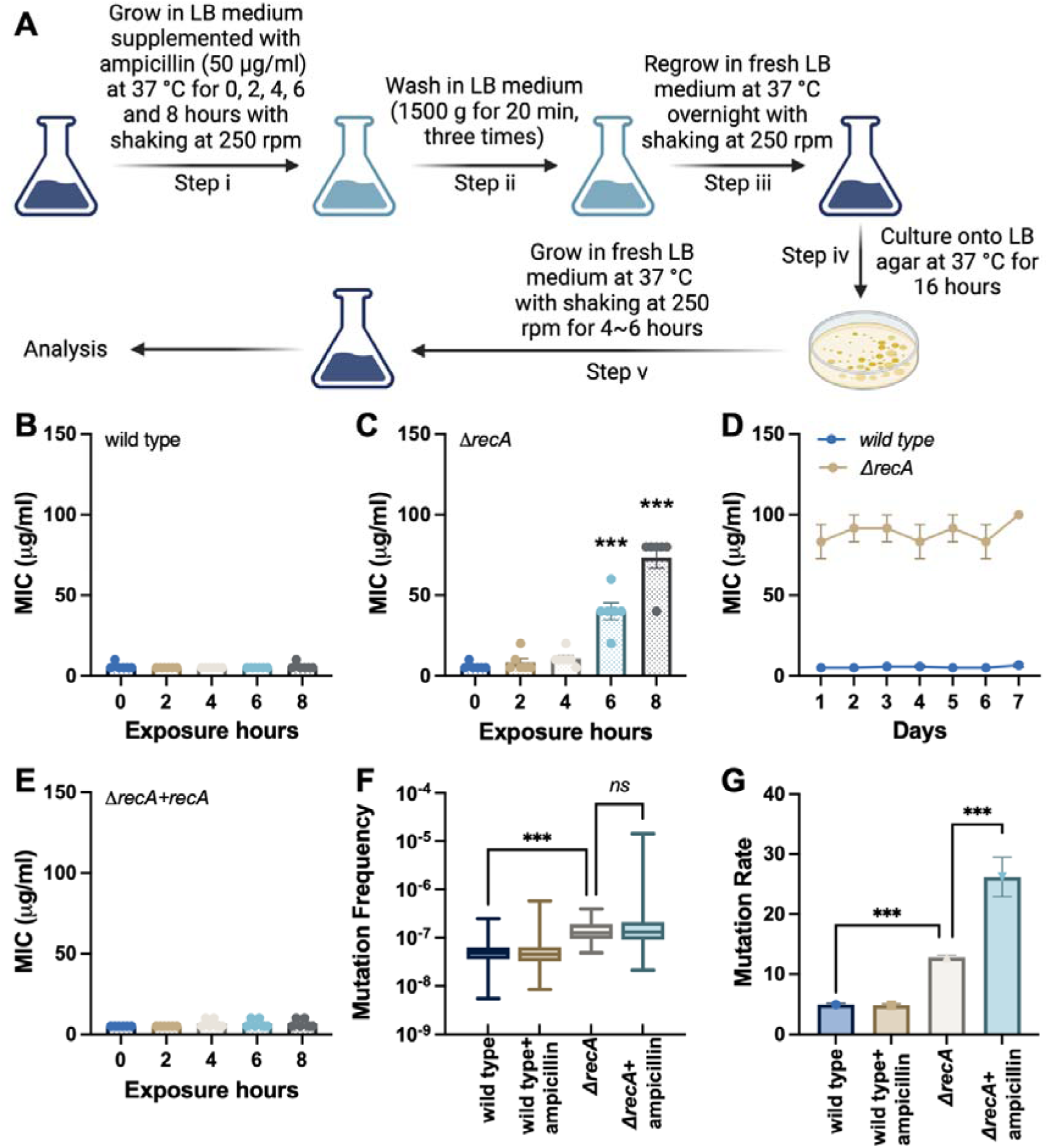
Fast evolution of antibiotic resistance in *E. coli recA* mutant strain. **(A)** Experimental flow for the single exposures to antibiotics in *E. coli* strains. Step i: an overnight culture (1 x 10^9^ CFU/mL cells) was diluted 1:50 into 30 mL LB medium supplemented with 50 μg/mL ampicillin and incubated at 37°C with shaking at 250 rpm for 0, 2, 4, 6 and 8 hours; Step ii: after each treatment, the ampicillin containing medium was removed by washing twice in a fresh LB medium; Step iii: the surviving isolates were resuspended in 30 mL fresh LB medium and regrown overnight at 37°C with shaking at 250 rpm; Step iv: cell cultures were plated onto LB agar and incubated for 16 hours at 37°C; Step v: single colonies were inoculated in 30 mL fresh LB medium and cultured at 37°C with shaking at 250 rpm for 4 to 6 hours. **(B)** MICs of ampicillin were measured against the wild type *E coli* strain after single exposures to ampicillin. **(C)** MICs of ampicillin were measured against the Δ*recA* strain after single exposures to ampicillin. **(D)** After the treatment of Step v, cells were continuously cultured in an antibiotic-free medium for seven days. MICs of ampicillin were measured each day. **(E)** MICs of ampicillin were measured against the Δ*recA* strain treated with ampicillin, where the expression of RecA was restored using plasmid-based constitutive expression of *recA* before the treatment of Step i. **(F)** Distribution of rifampicin-resistant mutant counts following single β-lactam exposure (96 replicate cultures in each group). Statistical comparison of mutation frequency (median values) used the Kruskal-Wallis test followed by Dunn’s multiple comparisons. **(G)** Mutation rate (mutations per culture) estimates derived by maximum likelihood analysis. Each experiment was independently repeated at least six times using parallel replicates, and the data are shown as mean ± SEM. Significant differences among different treatment groups are analysed by independent t-test, \**P* < 0.05, \*\**P* < 0.01, \*\*\**P* < 0.001, *ns*, no significance.

To assess the stability of this accelerated antibiotic resistance acquired by the Δ*recA* strain, we conducted a study wherein the Δ*recA* resistant isolates, originating from the initial 8-hour treatment with ampicillin, were continuously cultivated in a medium devoid of antibiotics for a period of seven days. Our findings revealed that once resistance was established, resistance remained stable and was able to be passed on to subsequent generations even in the absence of ampicillin (Fig. 1D). Moreover, we performed a complementation experiment by introducing a plasmid containing *recA* under its native promoter into the Δ*recA* strain prior to Step i in Fig. 1A, that is, before exposing the cells to ampicillin. Interestingly, this complemented strain displayed a comparable MIC to the isogenic wild type strain and maintained its sensitivity even after ampicillin treatment for up to 8 hours (Fig. 1E).

Antibiotic resistance evolves through the combined effects of mutational events and selection imposed by antimicrobial pressure (28). To clarify which mechanism underlies the rapid emergence of resistance in the Δ*recA* strain, we systematically analysed the mutation frequency and distribution patterns in response to ampicillin treatment. Fig. 1F shows the distribution of rifampicin-resistant colony-forming units (CFUs) across 96 independent cultures of the wild type and Δ*recA* strains, with and without exposure to ampicillin (50 µg/mL for 8 hours). Although the treatment of ampicillin in the Δ*recA* strain displayed a highly skewed distribution with apparent jackpot cultures, non-parametric statistical comparison (Kruskal-Wallis with Dunn’s test) did not detect a significant difference in overall mutation frequency compared to untreated Δ*recA*. This suggests that while the median mutation burden remained stable, rare resistant outliers were significantly enriched, which was a pattern consistent with selection rather than broad mutagenesis (29). Further, using the maximum likelihood estimation (MLE) (30), we calculated the mutation rates (mutations per culture) for each group (Fig. 1G). While the Δ*recA* group exhibited a modest increase in baseline mutation rate compared to the wild type strain, the addition of ampicillin led to a significant increase in estimated mutation rate.

To further assess the statistical properties of these distributions, we fitted the observed data to a Poisson model (Figure 1-figure supplement 5A). The mutation frequency distributions in the wild type strain with or without the treatment of ampicillin conformed well to Poisson expectations, consistent with random spontaneous mutation events. In contrast, the single exposure to ampicillin significantly deviated from the Poisson model, indicating a non-random clonal enrichment process in the Δ*recA* strain (31). Finally, we applied a fluctuation test-based inference framework grounded in the Luria-Delbrück model to distinguish between mutation induction and clonal selection (32). As shown in Figure 1-figure supplement 5B, only the Δ*recA* strain treated with ampicillin exhibited a markedly non-Poisson distribution of mutant counts, characterized by a long right-skewed tail and the emergence of jackpot cultures. This distribution is inconsistent with uniform mutagenesis and instead supports a model in which antibiotic treatment selectively enriches rare early-arising resistant subpopulations. Together, these findings demonstrate that a single exposure to β-lactam antibiotics promotes a rapid and heritable evolution of antibiotic resistance in the Δ*recA* strain, predominantly through selection rather than *de novo* induction of mutations.

### Detection of drug resistance associated DNA mutations in *recA* mutant resistant isolates

In bacteria, resistance to most antibiotics requires the accumulation of drug resistance associated DNA mutations that can arise stochastically and, under stress conditions, become enriched through selection over time to confer high levels of resistance (33). Having observed a non-random and right-skewed distribution of mutation frequencies in Δ*recA* isolates following ampicillin exposure, we next sought to determine whether specific resistance-conferring mutations were enriched in Δ*recA* isolates following antibiotic exposure. We thereby randomly selected 15 colonies on non-selected LB agar plates from the wild type surviving isolates, and antibiotic screening plates containing 50 µg/mL ampicillin from the Δ*recA* resistant isolates, respectively, and performed whole-genome sequencing. We found that drug resistance associated mutations were present in all resistant isolates, including the mutations in the promoter of the β-lactamase *ampC (P_ampC_)* in 8 isolates, the ampicillin-binding target PBP3 (*ftsI*) in 1 isolate, and the AcrAB-TolC subunit AcrB (*acrB*) mutations in 6 isolates (Fig. 2A). A mutation in gene *nudG* was detected in wild type surviving isolates after the single exposure to ampicillin (Fig. 2A), which is involved in pyrimidine (d)NTP hydrolysis to avoid DNA damage (34). Other mutations were listed in the Table S1.

**Figure 2.**
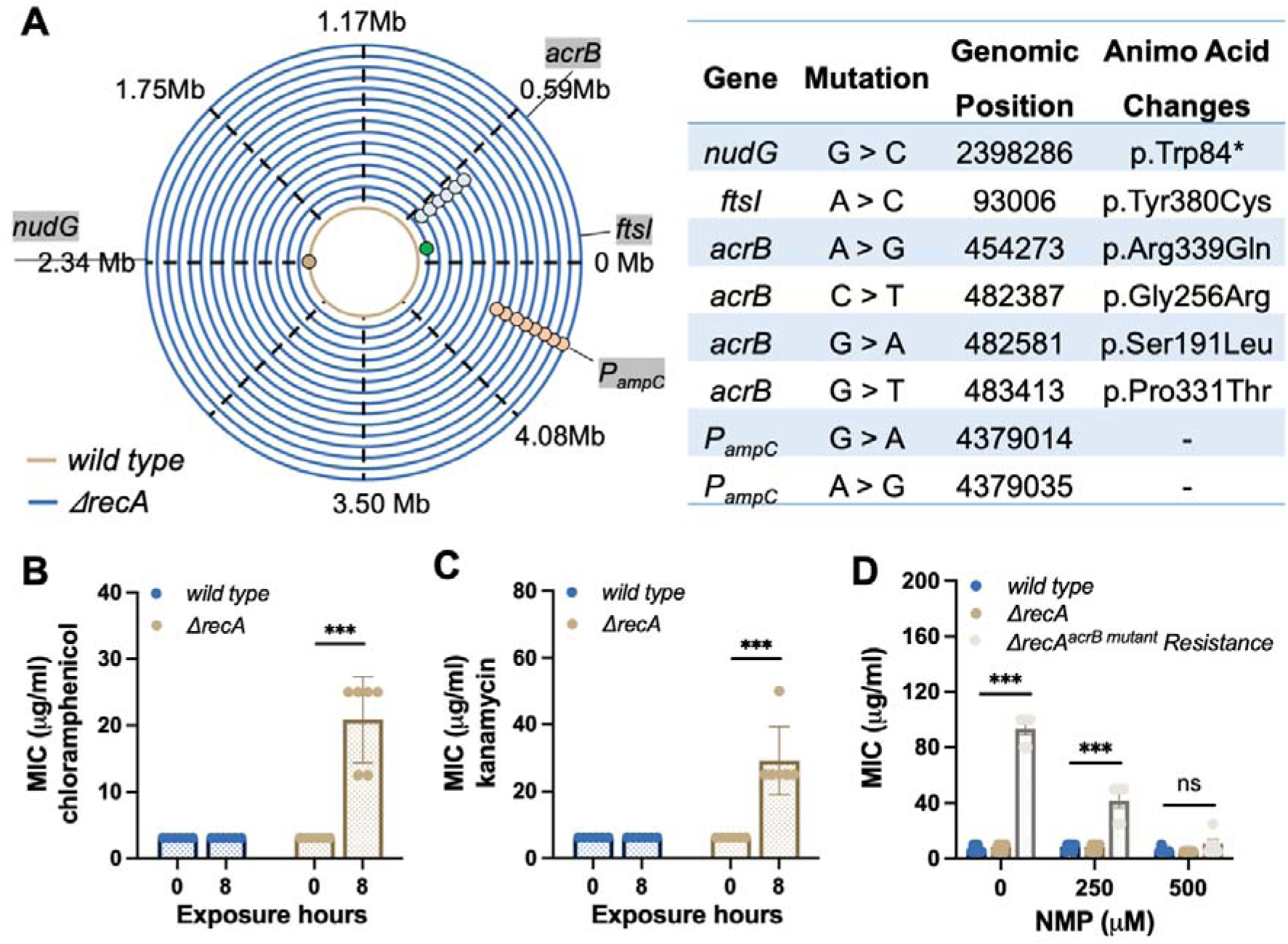
Rapid induction of drug resistance associated DNA mutations in the *recA* mutant strain. **(A)** Detection of drug resistance associated DNA mutations in the wild type and Δ*recA* strain after the single exposures to ampicillin at 50 µg/mL for 8 hours. **(B)** MICs of chloramphenicol against the wild type and Δ*recA* strain after the single treatment with ampicillin were tested. **(C)** MICs of kanamycin against the wild type and Δ*recA* strain after the single treatment with ampicillin were tested. **(D)** The wild type, Δ*recA*, and Δ*recA^acrB^ ^mutant^* resistant isolates were incubated with NMP at different concentrations for 12 hours. Subsequently, MICs were tested in these strains for resistance to ampicillin. Each experiment was independently repeated at least six times, and the data is shown as mean ± SEM. Significant differences among different treatment groups are analyzed by independent t-test, \**P* < 0.05, \*\**P* < 0.01, \*\*\**P* < 0.001, *ns*, no significance.

The presence of *P_ampC_* mutations was accompanied by a significant increase in the production of β-lactamase in bacteria (Figure 2-figure supplement 1). This leads to specific resistance to β-lactam antibiotics. The gene *acrB* codes for a sub-component of the AcrAB-TolC multi-drug efflux pump, which is central in Gram-negative bacteria (35,36). Mutations in AcrAB-TolC enhance the efflux of antibiotics and confer resistance to multiple drugs (35). Consequently, after short-term exposure to ampicillin, the Δ*recA* isolates carrying the *acrB* mutations exhibited resistance to other types of antibiotics, such as chloramphenicol and kanamycin (Fig. 2B and C). Treatment with high concentrations of 1-(1-Naphthylmethyl) piperazine (NMP), an efflux pump inhibitor (EPI) that competitively blocks TolC-composed efflux pumps, successfully restored the sensitivity of Δ*recA* resistant isolates to ampicillin, bringing them back to the equivalent concentrations found in the wild type (Fig. 2D). Collectively, these results suggest that β-lactam treatment rapidly selects for resistance-conferring mutations, which were enriched in Δ*recA* isolates following short-term exposure.

### Hindrance of SOS-independent DNA repair in *recA* mutant resistant isolates

Impairment of DNA damage repair can accelerate the accumulation of mutations and influence bacterial adaptability under antibiotic stress. While RecA is best known for regulating the SOS response, we asked whether its absence also impacts resistance evolution through SOS-dependent or -independent repair mechanisms. To investigate it, we first tested the ability of various mutants involved in different pathways of the SOS response to evolving antibiotic resistance following a single treatment with ampicillin for 8 hours at 50 µg/mL. A mutant form of the SOS master regulator LexA (*lexA3*), which is incapable of being cleaved and thus defective in SOS induction, did not exhibit antibiotic resistance evolution (Fig. 3A). Additionally, the deletion of either DpiB or DpiA (encoded by *citB* or *citA*, respectively), inhibiting the DpiBA two-component signal transduction system, did not result in the development of antibiotic resistance after ampicillin exposure (Fig. 3A). Moreover, the deletion of several downstream effectors of the SOS response, including those involved in cell division inhibition (SulA and YmfM encoded by *sulA* and *ymfM*) (37) (Fig. 3A), and DNA repair (DNA Pol II, DNA Pol IV, and DNA Pol V encoded by *polB*, *dinB* and *umuDC*) (38) also did not lead to the evolution of antibiotic resistance (Fig. 3A). These results indicate that the rapid resistance evolution observed in the Δ*recA* strain is not mediated by SOS pathway mutants, suggesting that RecA governs resistance evolution through SOS-independent mechanisms.

**Figure 3.**
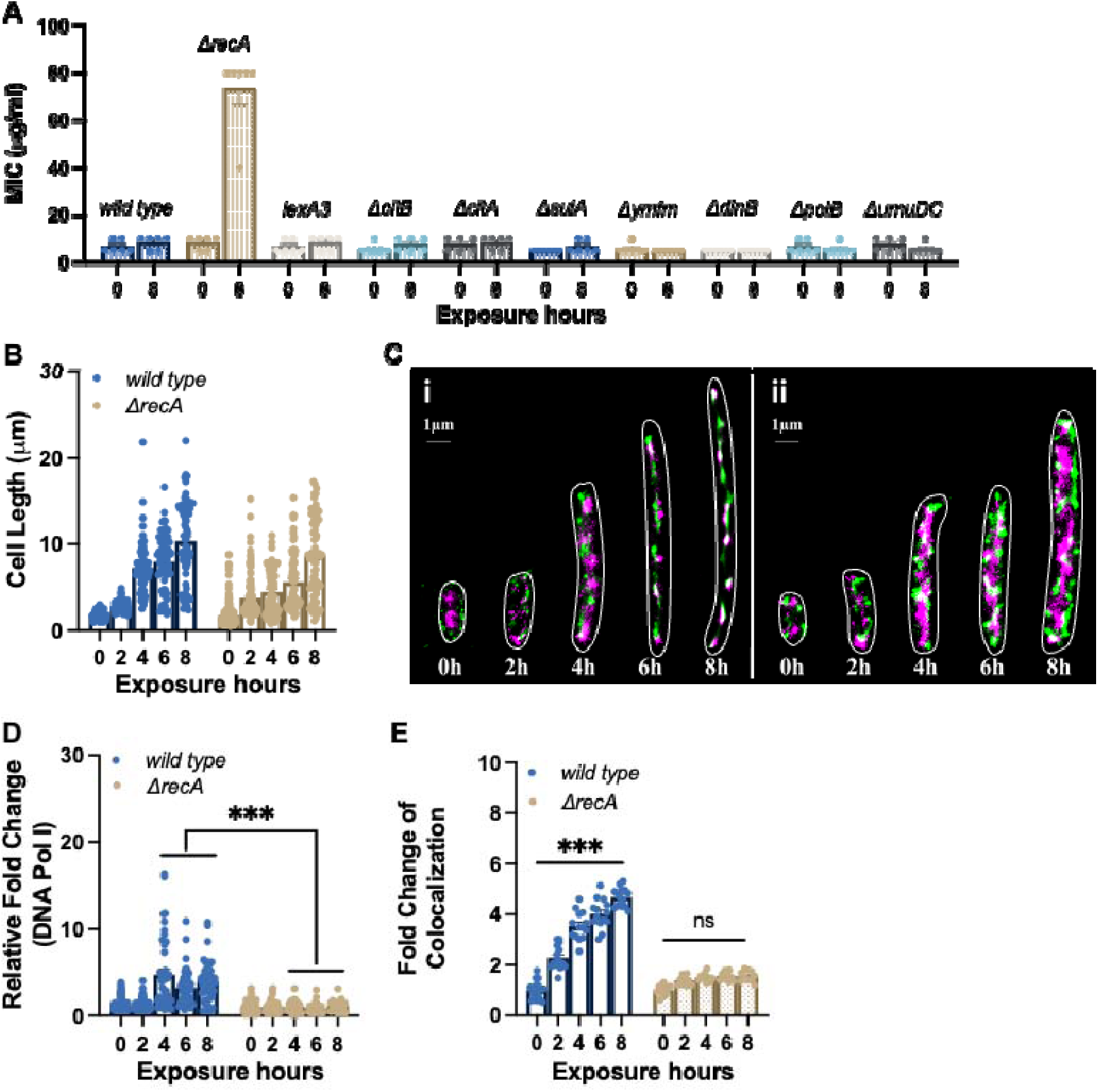
SOS-independent impairment of DNA repair in Δ*recA* resistant isolates. **(A)** MICs of ampicillin were tested against the wild type strain, Δ*recA* strain, and mutants lacking specific genes from the SOS response after single exposures to ampicillin at 50 µg/mL for 8 hours. **(B)** Filament cell lengths in the wild type (*n=253*) and the Δ*recA* strain (*n=216*) after single treatments with ampicillin at 50 μg/mL. **(C)** Multinucleated filaments were observed in the wild type (i) and the Δ*recA* (ii) strain after single exposures to ampicillin at 50 μg/mL. Purple: *E. coli* chromosome; green: DNA Pol I. **(D)** Relative fold changes of DNA Pol I in the wild type and Δ*recA* strain after single treatments with ampicillin at 50 μg/mL. **(E)** Co-localization between the *E. coli* chromosome and DNA Pol I in the wild type and Δ*recA* strain after the single exposures to ampicillin at 50 μg/mL. Data is shown as mean ± SEM. Significant differences among different treatment groups are analyzed by independent t-test, \**P* < 0.05, \*\**P* < 0.01, \*\*\**P* < 0.001, *ns*, no significance.

In addition to the DNA repair components associated with the SOS response, DNA Pol I plays a role in processing RNA primers during lagging-strand synthesis and filling small gaps during DNA repair reactions (39). Since DNA Pol I (encoded by *polA*) has been demonstrated as an essential gene required for the growth of *E. coli* in rich medium, including the LB medium (40,41), we next utilized Single Molecule Localization Microscopy (SMLM) to precisely locate the chromosome and DNA Pol I in a dynamic manner. During an 8 hour exposure to ampicillin, we observed the formation of multinucleated filaments in both the wild type and Δ*recA* strain, indicating a pause in cell division and suggesting a time period for bacterial DNA repair to take place (Fig. 3B and C) (42). However, the expression level of DNA Pol I was significantly suppressed in the Δ*recA* strain compared to the wild type strain after 4 hours of ampicillin exposure (Fig. 3D). More notably, the super-resolution colocalization analysis revealed a significantly lower ratio of co-localization between the chromosome and DNA Pol I in the Δ*recA* strain (Fig. 3E). Together, these findings demonstrate that the RecA loss impairs DNA repair capacity beyond SOS regulon. This repair deficiency contributes to genetic instability and is able to facilitate the rapid evolution of antibiotic resistance through SOS-independent pathways.

### Repression of antioxidative gene expression promotes the evolution of antibiotic resistance in the *recA* mutant strain

To further comprehend the fast evolution of β-lactam resistance observed in this study, we investigated the gene expression changes induced by ampicillin using a comprehensive transcriptome sequencing approach (total RNA-seq). Our analysis revealed significant transcriptomic alterations in both the wild type and Δ*recA* strain isolates following a single treatment with ampicillin, compared to untreated controls (Fig. 4A). Specifically, we identified changes in the expression of 161 and 248 coding sequences (with log_2_FC > 2 and *P* value < 0.05) in the wild type and Δ*recA* strains, respectively. Principal component analysis (PCA) demonstrated a notable disparity in the effects of ampicillin on the Δ*recA* strain compared to the wild type strain (Fig. 4B). Additionally, Venn diagram analysis confirmed that 138 and 225 genes were uniquely regulated by ampicillin exposure in the wild type and Δ*recA* strains, respectively (Fig. 4C).

**Figure 4.**
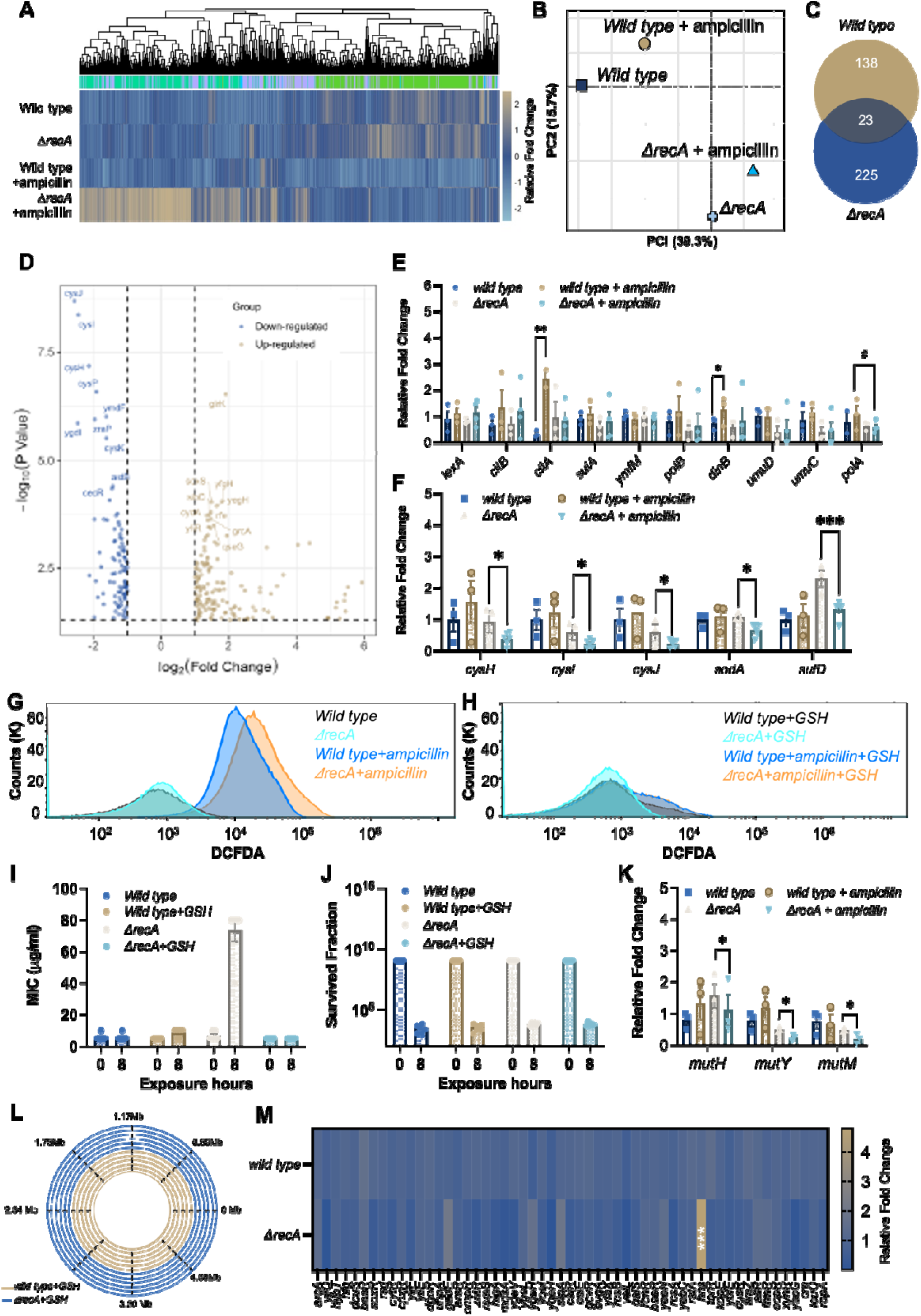
Overaccumulation of ROS drives the fast evolution of multi-drug resistance in the Δ*recA* strain. **(A)** Clustered heatmap of relative expression of coding sequences in the wild type and Δ*recA* strain with significant fold changes (log_2_FC > 2 and *P* value < 0.05). **(B)** Principal-component analysis (PCA) of normalised read counts for all strains. **(C)** Venn diagram of differentially expressed genes (log_2_FC > 2) after treatment with ampicillin at 50 μg/ml for 8 hours in the wild type and Δ*recA* strain. **(D)** The top 10 most differentially expressed genes in the Δ*recA* strain after the single treatment with ampicillin are labelled in each plot. Blue dots indicate genes with a significant downregulation compared to the untreated control (log_2_FC > 2 and *P* value < 0.05), and yellow dots indicate genes with a significant upregulation compared to the untreated control (log_2_FC > 2 and *P* value < 0.05). **(E)** Levels of transcription of SOS response system-associated genes and gene *polA* in the wild type and Δ*recA* strain after single exposures to ampicillin for 8 hours. **(F)** Levels of transcription of different antioxidative associated genes in the wild type and Δ*recA* strain after single exposures to ampicillin for 8 hours. **(G)** ROS levels were measured by flow cytometry before and after 8 hours of ampicillin treatment (50 μg/mL) in the wild type and Δ*recA* strains. **(H)** ROS levels were measured by flow cytometry in the wild type and Δ*recA* strains before and after 8 hours of ampicillin treatment at 50 μg/mL with the addition of GSH (50 mM). **(I)** The addition of 50 mM antioxidative compound GSH prevented the evolution of antibiotic resistance to ampicillin in the Δ*recA* strain treated with ampicillin at 50 μg/ml for 8 hours. **(J)** Survival fraction after a single exposure to ampicillin at 50 μg/ml for 8 hours in the wild type and the Δ*recA* strain with or without the addition of GSH at 50 mM. **(K)** Levels of transcription of proteins involved in the BER DNA repair system in the wild type and Δ*recA* strain after single exposures to ampicillin for 8 hours. **(L)** Whole genome sequencing confirms undetectable DNA mutations in the wild type and Δ*recA* strain treated with single exposures to ampicillin with the addition of GSH at 50 mM for 8 hours. **(M)** Transcription levels of all transcriptional repressors in the wild type and Δ*recA* strain after single treatments with ampicillin for 8 hours. Total RNA-seq was performed with three repeats in each group. Each experiment was independently repeated at least six times, and the data are shown as mean ± SEM. Significant differences among different treatment groups are analyzed by independent t-test, \**P* < 0.05, \*\**P* < 0.01, \*\*\**P* < 0.001, #*P* < 0.05.

To elucidate the differential expression of genes associated with specific biological functions, we conducted Gene Ontology (GO) enrichment analyses (Figure 4-figure supplement 1A and B) and Kyoto Encyclopedia of Genes and Genomes (KEGG) pathway analyses (Figure 4-figure supplement 1C and D). Our findings indicate that ampicillin profoundly impacted persistence pathways in the wild type strain, specifically affecting pathways related to quorum sensing, flagellar assembly, biofilm formation, and bacterial chemotaxis (43,44). Conversely, in the Δ*recA* strain, a distinct functional category associated with the oxidative stress response exhibited significant and unique down-regulation. This category included activities such as sulfate transporter activity, iron-sulfur cluster assembly, oxidoreductase activity, and carboxylate reductase activity.

To identify specific genes showing significant fold changes (log_2_FC > 2 and *P* value < 0.05), we used volcano plots to visualize the comprehensive changes in gene expression across the genome (Fig. 4D). We examined the transcription levels associated with the SOS response system and found that the transcription of several proteins in the wild type strain can be significantly induced by the single exposure to ampicillin, including *citB* and *dinB* (Fig. 4E). However, in the *recA* mutant strain, antibiotic exposure does not affect the transcription levels of any SOS system-related proteins, suggesting that antibiotic exposure induced the SOS response in the wild type strain but not in the Δ*recA* strain. More importantly, we discovered that the induction of the transcription level of the DNA Pol I was significantly suppressed after the single treatment of ampicillin in the Δ*recA* strain compared with that in the wild type strain (Fig. 4E). This is consistent with our imaging results and further supports the notion that an SOS-independent evolutionary mechanism dominates the development of antibiotic resistance in *recA* mutant *E. coli*.

Further, significant downregulation in the transcription of antioxidative-related genes in the Δ*recA* strain was detected, including *cysJ*, *cysI*, *cysH*, *soda*, and *sufD* (Fig. 4F). This downregulation suggested an excessive accumulation of reactive oxygen species (ROS) due to compromised cell antioxidative defences. It has been previously reported that the induction of mutagenesis can be stimulated by the overproduction of ROS during antibiotic administration, leading to the evolution of antibiotic resistance both *in vivo* and *in vitro* (22,23). Therefore, we hypothesized that elevated ROS levels facilitate the rapid evolution of antibiotic resistance in the Δ*recA* strain by increasing genetic instability and enabling the selection of resistant variants. To test it, we first examined the level of ROS generation in the wild type and Δ*recA* strains treated with ampicillin for 8 hours by using the fluorescent probe DCFDA/H2DCFDA. We found that ROS levels significantly increased in both the wild type and Δ*recA* strain after 8 hours of ampicillin treatment. However, ROS levels in the Δ*recA* strain showed a significant further increase compared to the wild type strain (Fig. 4G). Additionally, with the addition of 50 mM glutathione (GSH), a natural antioxidative compound, no significant change in ROS levels was observed in either the wild type or Δ*recA* strain before and after ampicillin treatment (Fig. 4H). Further, we supplemented the wild type and Δ*recA* strains with 50 mM GSH and treated them with ampicillin at a concentration of 50 μg/ml for 8 hours. Remarkably, the addition of GSH prevented the development of resistance to ampicillin in the Δ*recA* strain (Fig. 4I), without impairing the bactericidal effectiveness of ampicillin (Fig. 4J).

Apart from the SOS response, bacterial cells coordinate other DNA repair activities through a network of regulatory pathways, including base excision repair (BER) (45–47). The excessive generation of ROS results in elevated levels of deoxy-8-oxo-guanosine triphosphate (8-oxo-dGTP), an oxidized form of dGTP that becomes both highly toxic and mutagenic upon integration into DNA. The presence of 8-oxo-dG can induce SNP mutations, especially those occurring in guanine, which can be actively rectified by the BER repair pathway (47). Notably, BER glycosylases MutH and MutY can identify and repair these 8-oxo-dG-dependent mutations; however, when MutY and MutH are inactivated, unrepaired 8-oxo-dG can lead to the accumulation of SNP mutations within cells (47). As a result, we conducted a further assessment of the transcription levels of the BER repair pathway in both the wild type and Δ*recA* strain before and after a single exposure to ampicillin. We discovered that after an 8-hour treatment of ampicillin, three DNA repair-associated proteins, including MutH, MutY, and MutM, were notably suppressed in the Δ*recA* strain (Fig. 4K).

Finally, we sequenced the surviving Δ*recA* isolates and found that the addition of GSH inhibited drug resistance associated mutations in the Δ*recA* strain, which was detected in the Δ*recA* resistant isolates including genes of the promoter of *ampC* and *acrB* (Fig. 4L). Given the DNA repair impairment resulting in the generation of ROS, it would have been expected for genes involved in the oxidative stress response to be induced in RecA-deficient cells. However, the repression of antioxidative-related genes indicated the involvement of transcriptional repressors that might be regulated by RecA. Consequently, we examined the transcription levels of all transcriptional repressors in both the wild type and Δ*recA* strain. Remarkably, we observed a significant upregulation of H-NS, a crucial transcriptional repressor (Fig. 4M). This finding suggests that the upregulation of H-NS could potentially contribute to the suppression of antioxidant gene expression in the Δ*recA* strain, thereby promoting ROS accumulation and subsequent resistance evolution. Together, these findings demonstrate that RecA deficiency not only impairs DNA repair but also suppresses the oxidative stress response, leading to elevated ROS and increased mutational load. This oxidative-genetic imbalance forms the basis for enhanced mutational supply in RecA-deficient cells

## Discussion

In this study, we challenge the prevailing view that disabling RecA and thereby inhibiting the SOS response can prevent bacteria from developing antibiotic resistance (10). While the SOS response and RecA have been extensively studied for their roles in antibiotic resistance evolution (48), we observed a remarkably fast and stable evolution of multidrug resistance in the *E. coli* Δ*recA* strains following a single exposure to β-lactam antibiotics. This phenomenon cannot be explained by canonical SOS-driven mutagenesis (49), but instead reflects the interplay of two distinct evolution forces: RecA loss increases mutational supply through DNA repair suppression and ROS accumulation, while antibiotic-induced lethal stress provides a selective environment that promotes the expansion of rare and resistance-conferring variants (Fig. 5).

**Figure 5.**
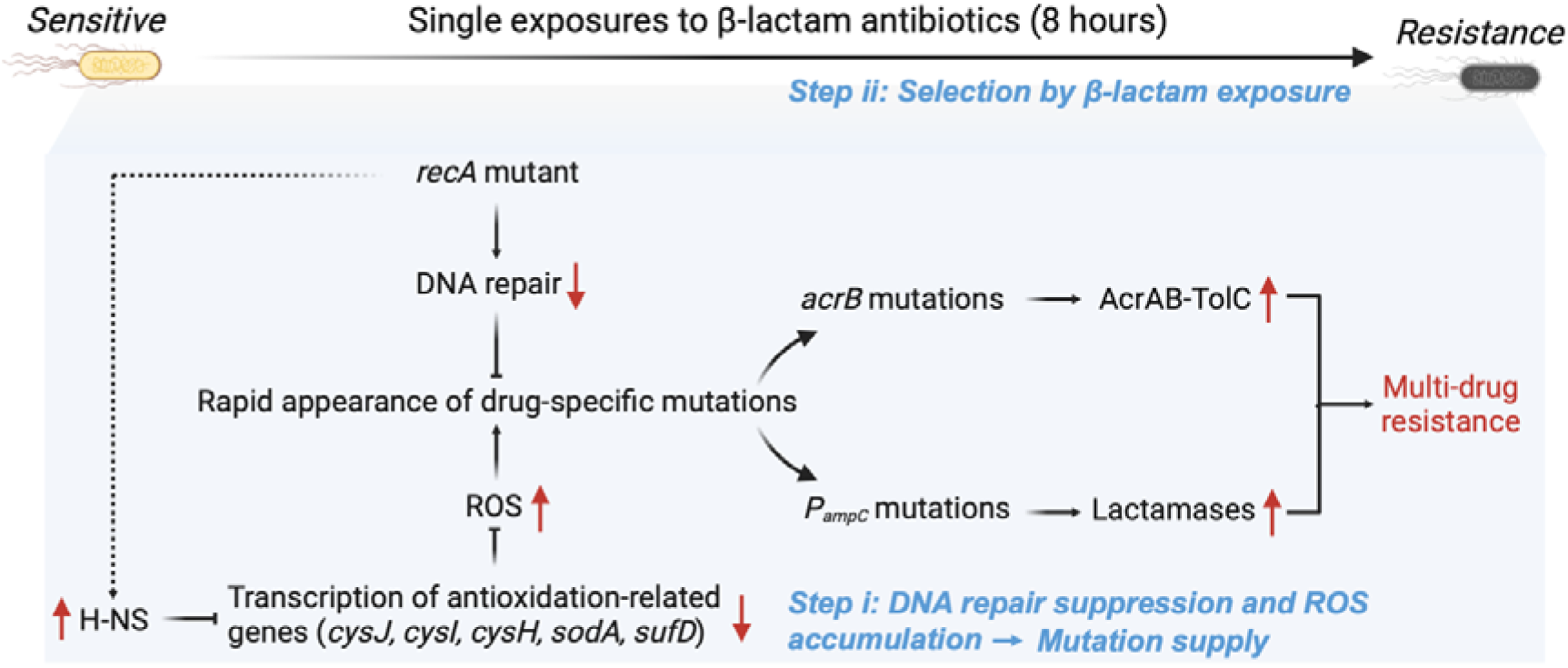
Mechanism of rapid development of multidrug resistance in the *recA* mutant *E. coli* strain. Deletion of *recA* impairs DNA damage repair and upregulates the transcription of the global repressor H-NS, though the mechanism of this regulation remains unclear. Elevated H-NS levels repress the expression of multiple antioxidant-related genes (*cysI*, *cysJ*, *cysH*, *sodA*, *sufD*), leading to excessive accumulation of ROS. The resulting oxidative stress increases the overall mutational burden. Upon single exposure to β-lactam antibiotics, drug-resistant subpopulations carrying specific mutations, such as in *acrB* or *P_ampC_*, are selectively enriched, ultimately driving the rapid emergence of multidrug resistance. This process reflects a two-step mechanism involving enhanced mutational supply and subsequent selection under antibiotic pressure.

To determine whether antibiotic exposure in Δ*recA* cells directly induces mutations or selectively enriches resistant variants, we combined statistical modelling, fluctuation analysis, and whole-genome sequencing. Although mutation rates estimated by maximum likelihood were moderately elevated in Δ*recA* cells after ampicillin treatment, the mutation frequency distributions were highly right-skewed and deviated from Poisson expectations, a hallmark of clonal selection rather than uniform and population-wide mutagenesis. These findings align with the classical Luria-Delbrück framework and indicate that resistance evolution in Δ*recA* is primarily driven by selection (29). However, our data also demonstrate that the mutational supply is enhanced in this background due to antibiotic-induced oxidative stress and impaired DNA repair, which together increase the likelihood that resistance-conferring mutations arise and persist. Thus, this process reflects a selection-dominated evolutionary trajectory facilitated by stress-enhanced mutagenesis, rather than classical induced mutation or selection on pre-existing variants.

Mechanistically, our data reveal that RecA plays broader roles in genome stability beyond its function in SOS activation. Disruption of individual SOS components, including *lexA*, *citA/B*, or translesion polymerases, did not recapitulate the Δ*recA*-specific resistance phenotype. Instead, Δ*recA* strains exhibited suppression of DNA Pol I. and key BER genes, such as *mutH*, *mutY*, and *mutM*, confirming that RecA is essential for the maintenance of repair fidelity under stress conditions. This defect likely enables ROS-induced DNA lesions to persist and become fixed as mutations.

In parallel, we identified a striking repression of antioxidative stress response genes in Δ*recA* strains exposed to ampicillin, including *cysJIH*, *sodA*, and *sufD*. This transcriptional suppression was associated with markedly elevated ROS levels. Crucially, supplementation with the antioxidant GSH reversed both ROS accumulation and the emergence of resistance, without impairing the bactericidal activity of ampicillin. This decouples the mutagenic and lethal effects of ROS, and highlights ROS as a driver of mutational supply rather than survival.

To explore the transcriptional basis for oxidative dysregulation, we examined global repressors and discovered that *hns* was significantly upregulated in Δ*recA* strains after ampicillin exposure. H-NS is a known transcriptional silencer of stress response genes, including those involved in redox regulation (50,51). We propose that RecA deficiency may lead to H-NS derepression, thereby silencing antioxidant defences, exacerbating ROS accumulation, and enabling mutation accumulation under stress. Although previous findings report that H-NS can down-regulate the transcriptional expression of *cysIJH* through the mediation of *cysB* (52,53), the mechanistic relationship between RecA and H-NS regulation remains to be experimentally validated.

More broadly, our findings highlight the repair-redox axis as a central regulator of bacterial evolvability. Rather than solely targeting bacterial growth or survival, future antimicrobial strategies might focus on constraining mutational potential. For example, co-administration of antioxidants or repair stabilizers could buffer stress-induced mutagenesis without compromising antibiotic efficacy, which has been a concept already under exploration in cancer therapeutics (54,55).

Clinically, the emergence of *acrB* mutations and enhanced activity of the AcrAB-TolC efflux pump in Δ*recA* strains recapitulates known multidrug resistance pathways. These observations raise concerns for therapies combining DNA repair inhibitors with ROS-inducing antibiotics or anticancer drugs (56). Our data suggest that such combinations may inadvertently promote resistance evolution, particularly in immunocompromised patients or during chemotherapy.

Finally, ROS-driven mutagenesis and repair suppression are not unique to β-lactams. Antimicrobial technologies such as antimicrobial photodynamic therapy (aPDT) and cold atmospheric plasma (CAP) (57,58) also rely on oxidative stress. In RecA-deficient or stress-sensitized bacteria, these approaches may risk accelerating resistance evolution unless accompanied by safeguards that preserve genomic integrity. Thus, maintaining the transcription of genes involved in the oxidative stress defence or combining antibiotics represents a promising strategy to prevent ROS-driven mutagenesis and thereby limit the evolutionary emergence of resistance during antimicrobial therapy.

## Acknowledgements

This work was supported by the Australian Research Council (ARC grant no: APP1165135), Science and Technology Innovation Commission of Shenzhen (KQTD20170810110913065), Australia China Science and Research Fund Joint Research Centre for Point-of-Care Testing (ACSRF658277, SQ2017YFGH001190).

## Data and materials availability

All data are available in the main text or the supplementary materials.

## Materials and Methods

### Bacterial strains, medium and antibiotics

Bacterial strains and plasmids used in this work are described in Table S2 and Table S3. Luria-Bertani (LB) was used as broth or in agar plates. *E. coli* cells were grown in LB liquid medium or on LB agar (1.5% w/v) plates at 37°C, unless stated otherwise, antibiotics were supplemented, where appropriate. Antibiotic stock solutions were prepared by dissolving antibiotics in MilliQ filter sterilising, including ampicillin (50 mg/mL), penicillin G (100 mg/mL), carbenicillin (20 mg/mL), kanamycin (50 mg/mL) and tetracycline (10 mg/mL). Chloramphenicol stock solution was prepared in 95% EtOH (25 mg/mL). Antibiotic solutions were stored at -20°C (long-term) or 4°C (short-term).

### Treatment with antibiotics to induce evolutionary resistance

For the single exposure to antibiotic experiment, an overnight culture (0.6 mL; 1 x 10^9^ CFU/mL cells) was diluted 1:50 into 30 mL LB medium supplemented with antibiotics (50 μg/mL ampicillin, 1 mg/mL penicillin G, or 200 μg/mL carbenicillin) and incubated at 37°C with shaking at 250 rpm for 0, 2, 4, 6 and 8 hours, respectively. After each treatment, the antibiotic-containing medium was removed by washing twice (20 min centrifugation at 1500 g) in fresh LB medium (See Fig. 1A for method overview).

To test resistance, the surviving isolates were first resuspended in 30 mL LB medium and grown overnight at 37°C with shaking at 250 rpm. The regrown culture was then plated onto LB agar and incubated overnight at 37°C. Single colonies were isolated and grown in LB medium for 4-6 hours at 37°C with shaking at 250 rpm, which were then used to test the resistance or stored at -80°C for future use.

For the ALE antibiotic treatment experiments, an overnight culture (0.6 mL; 1 x 10^9^ CFU/mL cells) was diluted 1:50 into 30 mL LB medium supplemented with 50 μg/mL ampicillin and incubated at 37°C with shaking at 250 rpm for 4.5 hours. After treatment, the antibiotic-containing medium was removed by washing twice (20 min centrifugation at 1500 g) in a fresh LB medium. The remaining pellet was resuspended in 30 mL LB medium and grown overnight at 37°C with shaking at 250 rpm. Ampicillin treatment was applied to the regrown culture and repeated until resistance was established, as confirmed by MIC measurement.

### Antibiotic susceptibility testing

The susceptibility of *E. coli* cells to antibiotics was measured using minimum inhibitory concentration (MIC) testing (60). In brief, overnight cultures were diluted and incubated at 37°C for 4-6 hours with shanking at 250 rpm. Cells were then diluted 1:100 and incubated with increasing concentrations of antibiotics in the Synergy HT BioTek plate reader (BioTek Instruments Inc., USA) at 37°C for 16 hours. It was programmed to measure the OD hourly at 595 nm (Gen5 software, BioTek Instruments Inc., USA). The minimum inhibitory concentration was determined as the concentration of antibiotic where no visible growth was observed.

### Survival assays

Overnight cultures of *E. coli* were prepared from single colonies in LB medium and incubated at 37°C with shaking (250 rpm). The overnight cultures were diluted 1:50 in fresh LB medium containing 50 µg/mL ampicillin to initiate antibiotic treatment. Cultures were incubated at 37°C for the indicated times under shaking conditions. Following treatment, cells were collected by centrifugation (1500 g, 20 minutes), and the antibiotic-containing medium was removed by washing twice with fresh LB medium. Serial dilutions of the washed cultures were prepared, and 25 µL of each dilution was plated onto LB agar plates. Plates were incubated overnight at 37°C, and CFUs were counted the following day to evaluate bacterial survival rates.

### Mutation frequency and fluctuation analysis

Overnight cultures inoculated from single colonies in LB medium were diluted 1:1,000,000 and incubated at 37°C with shaking until the OD_600_ reached to 1∼1.3. This extreme dilution minimizes the presence of pre-existing stationary phase mutants and allows *de novo* mutation events to occur during exponential growth. For each biological condition, 96 independent parallel cultures were prepared to perform a fluctuation analysis. The total number of colony-forming units per ml (CFU/ml) was determined by plating on LB agar. To identify rifampicin-resistant mutants, the remaining culture volume was centrifuged and plated on LB agar containing rifampicin (100 μg/mL). LB plates were incubated for 24 hours at 37°C and selective plates were incubated for 48-72 hours at 37°C (61).

The mutation frequency was calculated as the ratio of CFU/ml on rifampicin-containing plates to CFU/ml on non-selective LB plates for each culture. Distributions of mutation frequencies across replicate cultures were plotted, and deviations from Poisson expectations were assessed by model fitting.

To estimate the mutation rate (mutations per culture), maximum likelihood estimation (MLE) was applied using the Ma-Sandri-Sarkar algorithm implemented in the FALCOR toolset (https://www.keshavsingh.org/protocols/FALCOR.html). The inferred mutation rate distributions were compared across treatment conditions, and non-Poisson distributions indicative of jackpot cultures were further analysed using fluctuation test-based inference following the Luria–Delbrück framework (29).

### Construction of *recA* deletion mutant

Lambda Red recombination was used to generate the gene *recA* deletion in the *E. coli* K-12 strain, followed by previously reported methods with modifications (62,63). Primers (*recA-FWD* and *recA-REV,* Table S4) were designed approximately 50 bp upstream and downstream to the gene *recA* on the chromosome to amplify the tetracycline cassette as well as the flanking DNA sequence needed for homologous recombination. Phusion polymerase (NEB) was used to amplify the DNA sequence (Table S4), and the reaction was cleaned up using a PureLink™ PCR purification kit (ThermoFisher Scientific) as per the manufacturer’s instructions. Electro-competent *E. coli* MG1655 containing the recombinase plasmid pKD46 was transformed with 50 ng of amplified DNA a 30°C. The transformation was plated onto LB agar plates containing 10 μg/mL tetracycline and incubated overnight at 37°C. PCR was used to confirm the insertion of the tetracycline resistance cassette at the correct site on the chromosome using primers upstream and downstream to the gene *recA*. The newly constructed mutant strains were cured of plasmid pKD46 by incubating LB streak plates at 42°C overnight. Loss of the plasmid was confirmed by lack of ampicillin sensitivity on LB agar plates. Mutant strains were made electro-competent, and 50 μL of cells were transformed with plasmid pCP20 and incubated on 100 μg/mL ampicillin plates at 30°C overnight. A few colonies were then restreaked onto LB plates and incubated overnight at 42°C. PCR products confirmed the loss of cassette and plasmid.

### β-lactamase assay

The amount of β-lactamase was measured using a β-lactamase Activity Assay Kit (Sigma-Aldrich, US). Briefly, cells were collected by centrifugation at 10,000 g for 10 min, and the pellet was resuspended with 5 µL of assay buffer per mg of sample. Then, 48 μL of the sample was mixed with 2 μL of nitrocefin. The β-Lactamase activity was monitored by measuring the absorbance at 490□nm for 30□min at 28°C. The level of β-lactamase was determined by the absorbance at OD_390_.

### Whole genome sequencing

Resistant clones were isolated by selection using LB agar plates with the supplementation of ampicillin at 50 μg/mL, and non-resistant clones were isolated from the LB agar plates without the supplementation of antibiotics. Chromosomal DNA was extracted and purified using the PureLink™ Genomic DNA mini kit following the manufacturer’s instructions (ThermoFisher Scientific). Whole genome sequencing (WGS) was conducted following the Nextera Flex library preparation kit process (Illumina). Briefly, genomic DNA was quantitatively assessed using Quant-iT picogreen dsDNA assay kit (Invitrogen, USA). The sample was normalised to the concentration of 1 ng/μL. 10 ng of DNA was used for library preparation. After tagmentation, the tagmented DNA was amplified using the facility’s custom-designed i7 or i5 barcodes, with 12 cycles of PCR. The quality control for the samples was done by sequencing a pool of samples using MiSeq V2 nano kit - 300 cycles. After library amplification, 3 μL of each library was pooled into a library pool. The pool was then cleaned up using SPRI beads following the Nextera Flex clean-up and size selection protocol. The pool was then sequenced using a MiSeq V2 nano kit (Illumina, USA). Based on the sequencing data generated, the read count for each sample was used to identify the failed libraries (i.e., libraries with less than 100 reads).

Moreover, libraries were pooled at different amounts based on the read count to ensure equal representation in the final pool. The final pool was sequenced on Illumina NovaSeq 6000 Xp S4 lane, 2 × 150 bp. WGS read quality was assessed using FASTQC (version 0.11.5) and trimmed using Trimmomatic (version 0.36) with default parameters and trimmed of adaptor sequences (TruSeq3 paired-ended). Reads were aligned to the *E. coli* MG1655 genome (http://bacteria.ensembl.org/Escherichia_coli_str_k_12_substr_mg1655_gca_000005845/Info/Index/, assembly ASM584v2) and then analysed variants following GATK Best Practices for Variant Discovery (HaplotypeCaller) (64). Further genome variant annotation was conducted using the software SnpEff (65).

### Global transcriptome sequencing

After ampicillin treatment for 0 and 8, surviving isolates were immediately washed and harvested for global transcriptome sequencing. Total RNA was extracted from the cell pellets using a PureLink RNA mini kit (Invitrogen) as per the manufacturer’s instructions. The global transcriptome sequencing was processed and analysed by Genewiz, Jiangsu, China. Primers used in this work are listed in Table S4. RNA-Seq read quality was assessed using FASTQC and trimmed using Trimmomatic with default parameters. Reads were aligned to the *E. coli* MG1655 genome (http://bacteria.ensembl.org/Escherichia_coli_str_k_12_substr_mg1655_gca_000005845/Info/Index/, assembly ASM584v2) and then counted using the RSubread aligner with default parameters (66). After mapping, differential expression analysis was carried out using strand-specific gene-wise quantification using the DESeq2 package (67). Further normalisation was conducted using RUVSeq and the RUV correction method, with *k*□=□1 to correct for batch effects, using replicate samples to estimate the factors of unwanted variation (68). Absolute counts were transformed into standard z-scores for each gene over all treatments, that is, absolute read for a gene minus mean read count for that gene over all samples and then divided by the standard deviation for all counts over all samples. Genes with an adjusted *P* value (*P*_adj_)□of ≤ 0.05 were considered differentially expressed. PseudoCAP analysis was conducted by calculating the percentage of genes in each classification that were differentially expressed (log_2_FC ≥ ±2, *P*_adj_□≤□0.05).

### ROS measurement

Intracellular ROS accumulation was measured using the Cellular ROS Assay Kit (Abcam, ab113851) according to the manufacturer’s protocol. Overnight cultures of both wild type and Δ*recA* strains were grown in LB medium. Cells were treated with 50 µg/mL ampicillin for 8 hours at 37°C with shaking (250 rpm). After the treatment, cells were washed twice with 1x buffer to remove residual antibiotics and debris. The collected cells were resuspended in 1x buffer and incubated with the fluorescent ROS probe DCFDA (2’,7’–dichlorofluorescin diacetate)/H2DCFDA at a final concentration of 10 µM for 30 minutes at 37°C in the dark to prevent probe degradation.

After incubation, 500 µL of the stained cells were immediately analysed using a CytoFLEX LX flow cytometer (Beckman Coulter). Fluorescence was detected in the FITC channel (excitation at 488 nm, emission at 525 nm). Data were analysed using FlowJo software (Version X, Tree Star Inc.), with fluorescence intensity serving as an indicator of intracellular ROS levels. Appropriate controls, including unstained cells and cells without ampicillin treatment, were included to ensure accurate ROS measurement.

### Single-molecule localisation imaging and data analysis

Single-molecule localisation imaging was performed on a custom-built Stochastic Optical Reconstruction Microscope (STORM) with an Olympus IX81 microscope frame, a 100x magnification NA 1.45 objective (Olympus) and an EMCCD camera (DU-897, Andor) as described previously (69–71). In summary, 35 mm cell culture dishes (0.17 mm No.1 coverglass) were cleaned with 1 M KOH for 30 minutes in an ultrasonic cleaning machine, followed by three washes with MilliQ water. The dishes were air-dried with high-purity nitrogen blowing and sterilised by UV exposure for 30 minutes. *E. coli* cells were fixed with NaPO4 (30 nM), formaldehyde (2.4%), and glutaraldehyde (0.04%) at room temperature for 15 minutes, followed by 45 minutes on ice. Samples were then centrifuged to collect the pellet cells, and the supernatant was discarded. Cell pellets were washed twice with phosphate-buffered saline (PBS), pH 7.4. Cells were resuspended in 200 μL of GTE buffer and kept on ice until 200 µL was placed onto the coverslip bottom of the cleaned 35 mm culture dish. To label the bacterial chromosome, a Click-iT EdU kit was used prior to fixation following the manufacturer’s instruction (ThermoFisher) and as described before. To label DNA polymerase I, fixed cells were blocked and permeabilised with blocking buffer (5% wt/vol bovine serum albumin (Sigma-Aldrich) and 0.5% vol/vol Triton X-100 in PBS) for 30 min and then incubated with 1 μg/mL primary antibody against DNA polymerase I (ab188424, Abcam) in blocking buffer for 60 min at room temperature. After washing with PBS three times, the cells were incubated with 2 μg/mL fluorescently labelled secondary antibody (Alexa 647, A20006, ThermoFisher) against the primary antibody in the blocking buffer for 40 min at room temperature. After washing with PBS three times, the cells were postfixed with 4% (wt/vol) paraformaldehyde in PBS for 10 min and stored in PBS before imaging. STORM image analysis, drift correction, image rendering, protein cluster identification and images presentation were performed using Insight342, custom-written Matlab (2012a, MathWorks) codes, SR-Tesseler (IINS, Interdisciplinary Institute for Neuroscience) (72), and Image J (National Institutes of Health).

### Statistical analysis

Statistical analysis was performed using GraphPad Prism v.9.0.0. All data are presented as individual values and mean or mean ± SEM. Unless otherwise specified, statistical comparisons were conducted using one-way ANOVA (for multiple groups) or unpaired two-tailed Student’s t-tests (for two-group comparisons), assuming a 95% confidence interval. A probability value of *P* < 0.05 was considered significant. Statistical significance is indicated in each figure. All remaining experiments were repeated independently, at least six with similar results.

## Data availability

Sequence data supporting this study’s findings have been deposited in the GEO repository with the GEO accession number GSE179434.

---

**Figure 1-figure supplement 1.**
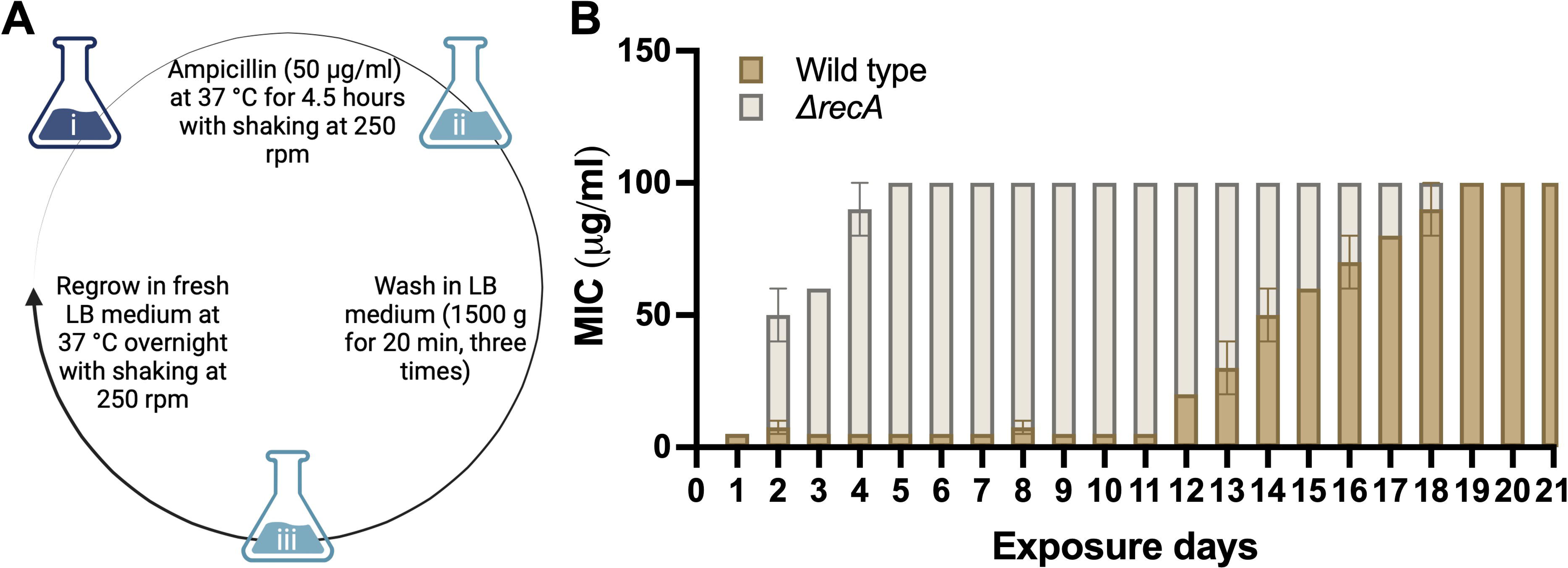
Long-term exposures to ampicillin induced the evolution of resistance in the wild type and Δ*recA E. coli* strain. **(A)** The experimental flow of ALE antibiotic treatment experiment. An overnight culture (0.6 mL; 1 x 10^9^ CFU/mL cells) was diluted 1:50 into 30 mL LB medium supplemented with 50 μg/mL ampicillin and incubated at 37°C with shaking at 250 rpm for 4.5 hours. After treatment, the antibiotic-containing medium was removed by washing twice (20 min centrifugation at 1500 g) in fresh LB medium. The surviving isolates were resuspended in 30 mL LB medium and grown overnight at 37°C with shaking at 250 rpm. Ampicillin treatment was applied to the regrown culture and repeated until resistance was established. **(B)** Changes in the MICs of ampicillin in the wild type and Δ*recA* strain after 21 days of treatment with ampicillin at 50 μg/ml for 4.5 hours each day. MICs were measured after each daily treatment. Each experiment was independently repeated at least twice, and the data are shown as mean ± SEM.

**Figure 1-figure supplement 2.**
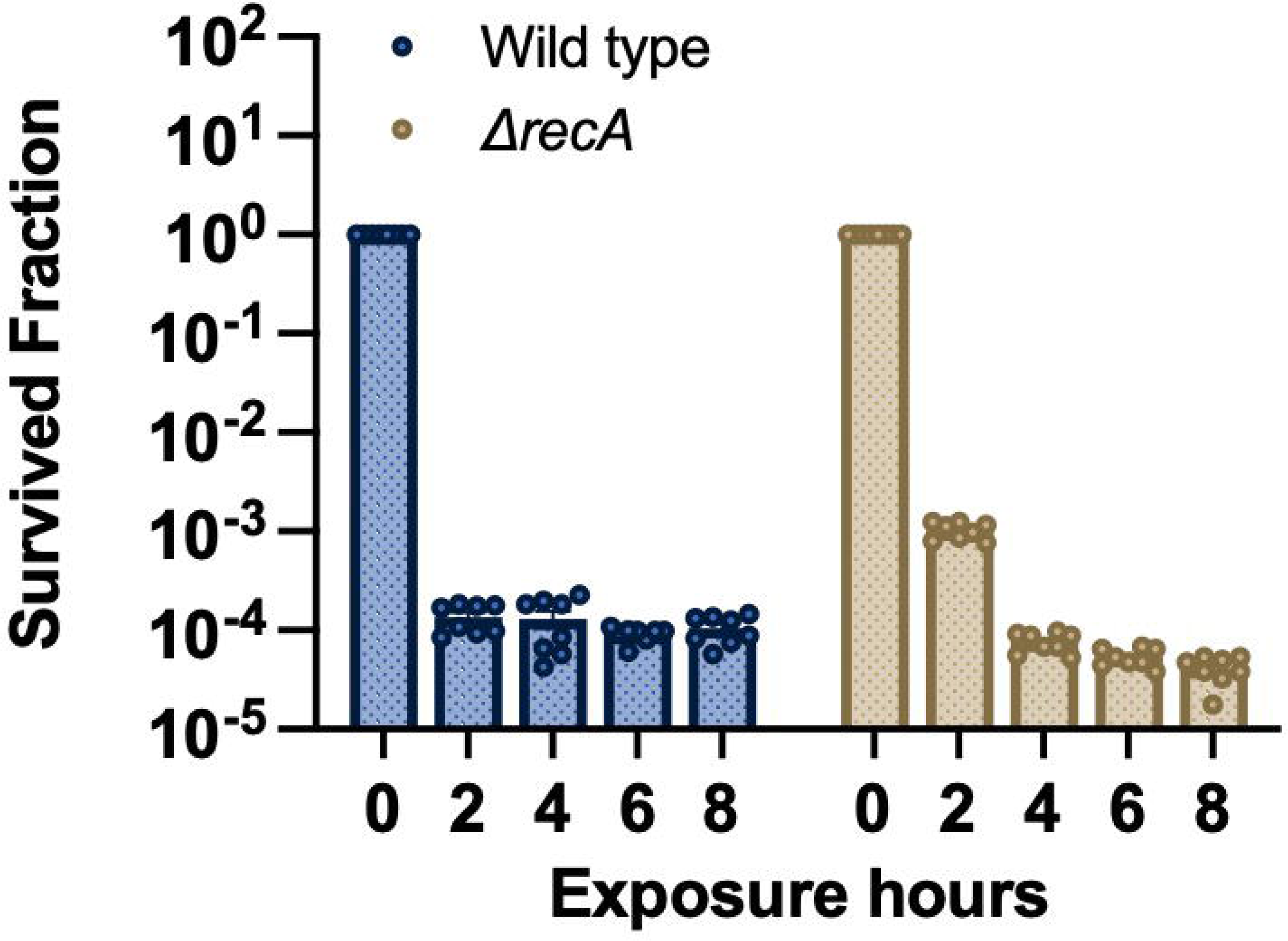
The survival rate after a single exposure to ampicillin at 50 μg/ml for 0,2,4,6, and 8 hours in the wild type and Δ*recA* strain.

**Figure 1-figure supplement 3.**
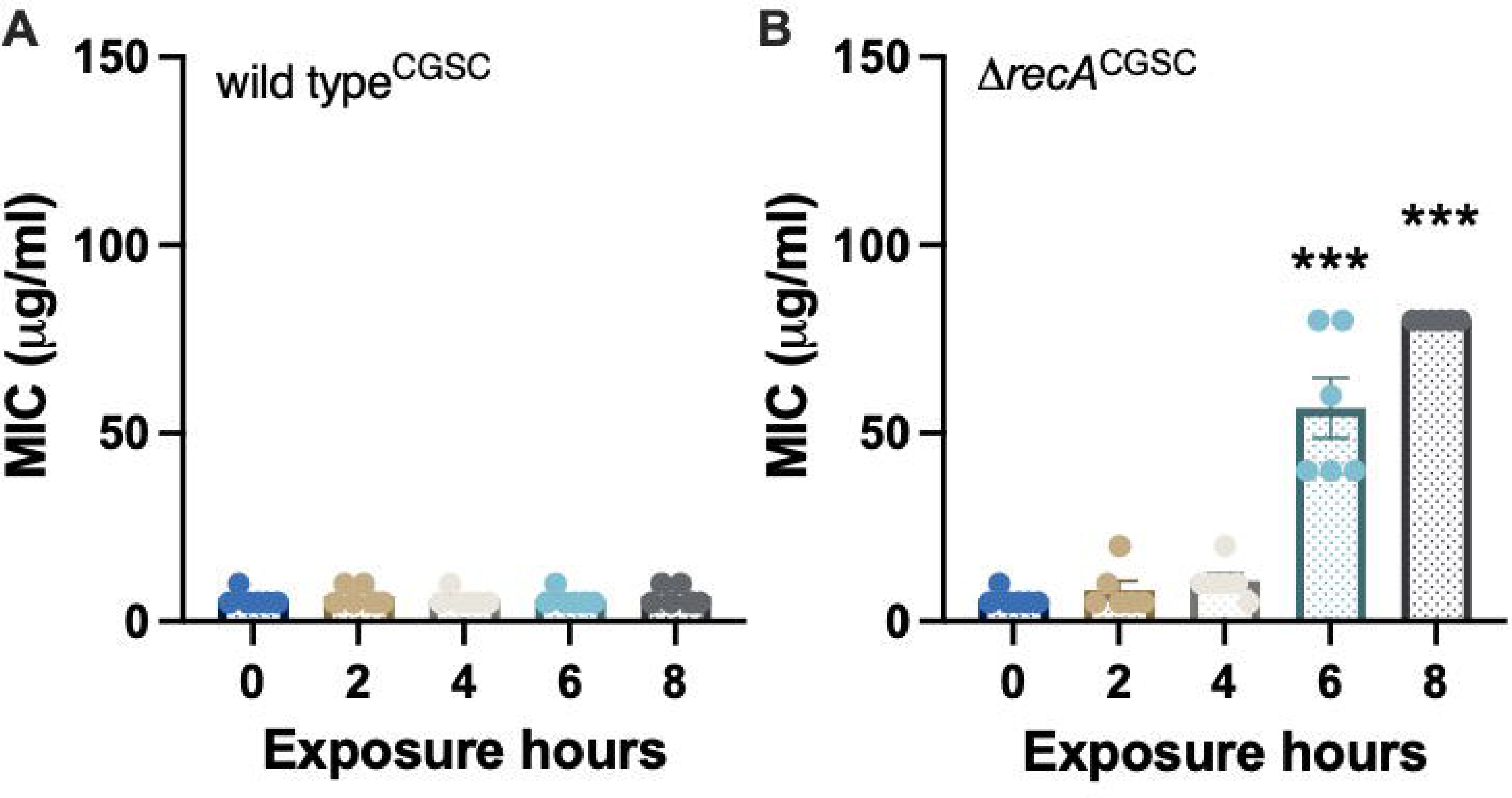
Single exposures to ampicillin induced the evolution of resistance in the Δ*recA^CGSC^*strain (JW2669-1). **(A)** MICs of ampicillin were measured against the wild type^CGSC^ *E coli* strain after single exposures to ampicillin. **(B)** MICs of ampicillin were measured against the Δ*recA^CGSC^* strain after single exposures to ampicillin. Each experiment was independently repeated at least six times using parallel replicates, and the data are shown as mean ± SEM. Significant differences among different treatment groups are analyzed by independent t-test, \**P* < 0.05, \*\**P* < 0.01, \*\*\**P* < 0.001.

**Figure 1-figure supplement 4.**
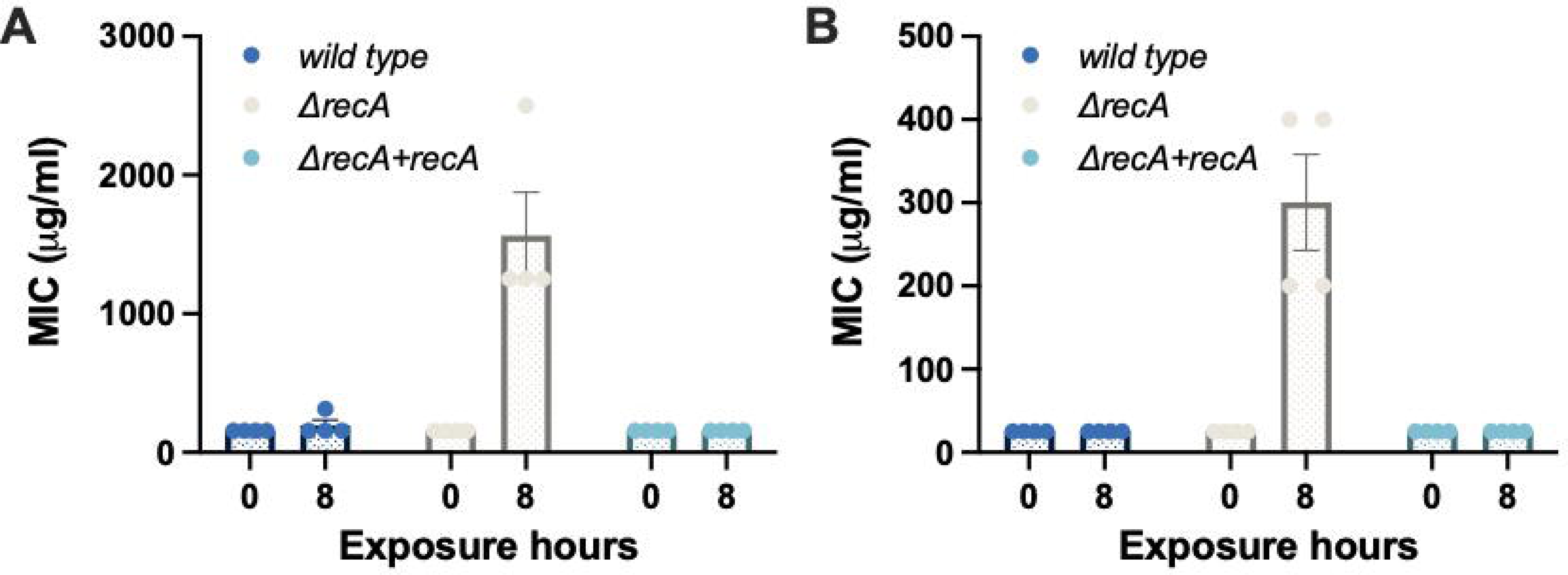
Single exposures to other β-lactam antibiotics induced the evolution of resistance in the Δ*recA* strain. **(A)** MICs of penicillin G in the wild type, Δ*recA*, and complemented Δ*recA* strain, where the expression of RecA was restored, after single exposures to penicillin G at 1 mg/mL for 8 hours. **(B)** MICs of carbenicillin in the wild type, Δ*recA*, and complemented Δ*recA* strain, where the expression of RecA was restored, after single exposures to carbenicillin at 200 µg/mL for 8 hours. Each experiment was independently repeated at least six, and the data are shown as mean ± SEM.

**Figure 1-figure supplement 5.**
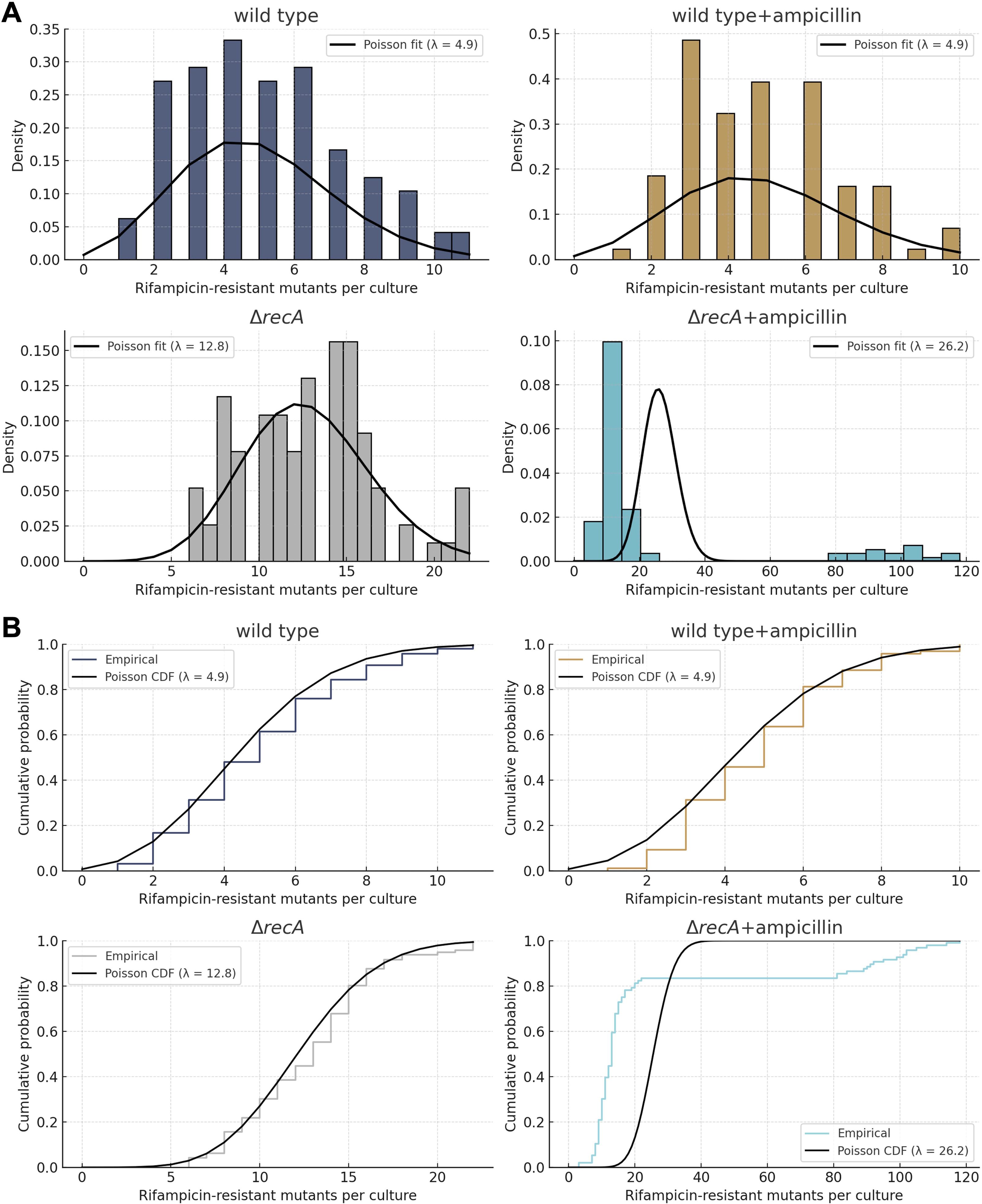
Distribution fitting and fluctuation analysis support a selection-driven resistance mechanism in the Δ*recA* strain. **(A)** Rifampicin-resistant mutant counts from replicate cultures were fitted to theoretical Poisson distributions based on observed means. **(B)** Fluctuation analysis using the Luria-Delbrück framework revealed long-tailed, highly skewed mutation count distributions in the Δ*recA* strain under the treatment of ampicillin, characteristic of jackpot cultures arising from early-arising resistant mutants.

**Figure 2-figure supplement 1.**
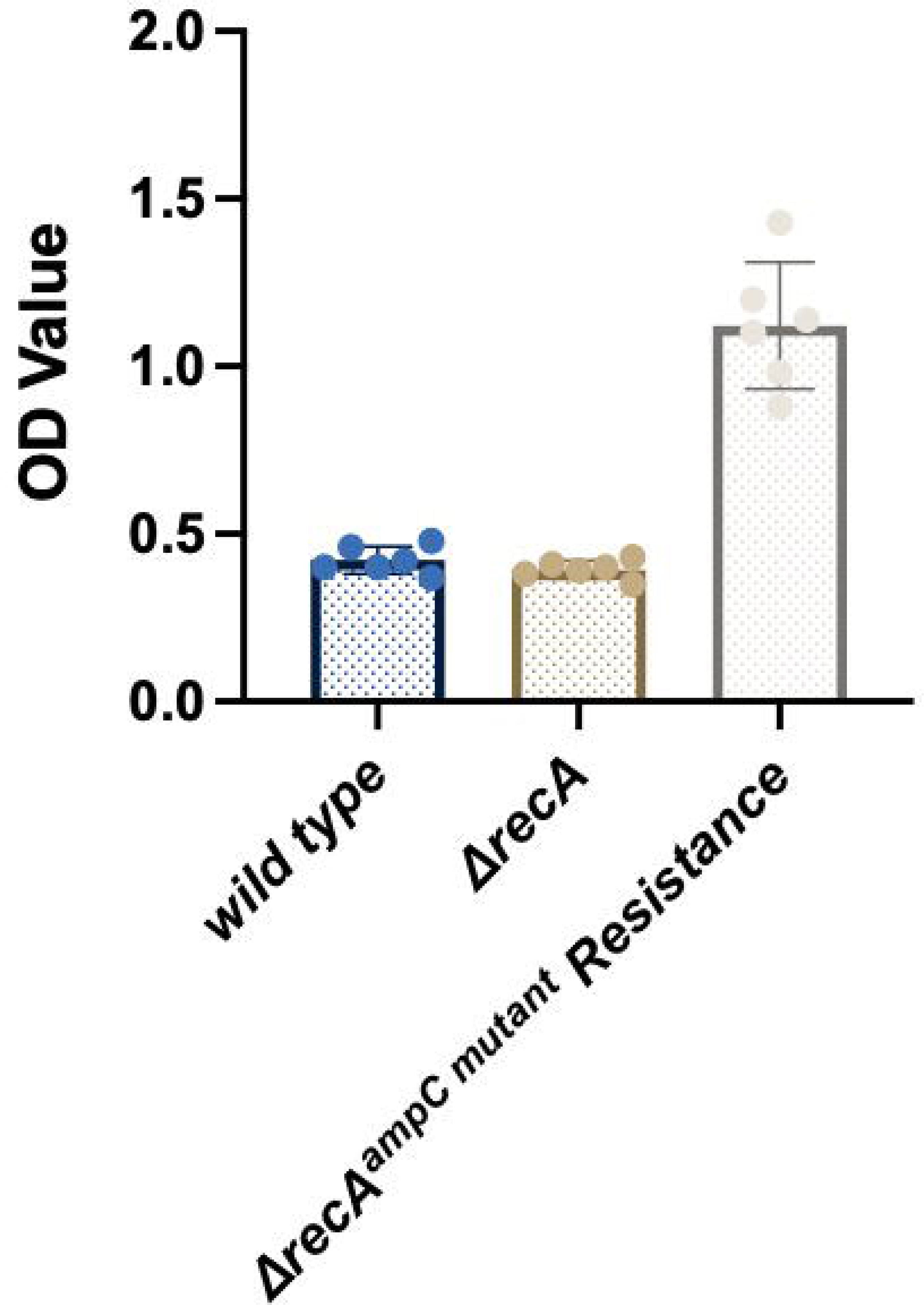
The activity of β-lactamase was increased in the Δ*recA* culture supernatants. The Δ*recA* strain was treated with ampicillin at 50 µg/ml for 8 hours, and surviving isolates harbouring the *ampC* mutations were selected. The level of β-lactamase in cell culture supernatants was determined by the absorbance at OD_490_. The levels of β-lactamase in the wild type or Δ*recA* strain culture supernatants without exposure to ampicillin were tested as a control. Each experiment was independently repeated six times. Each experiment was independently repeated at least six, and the data are shown as mean ± SEM.

**Figure 4-figure supplement 1.**
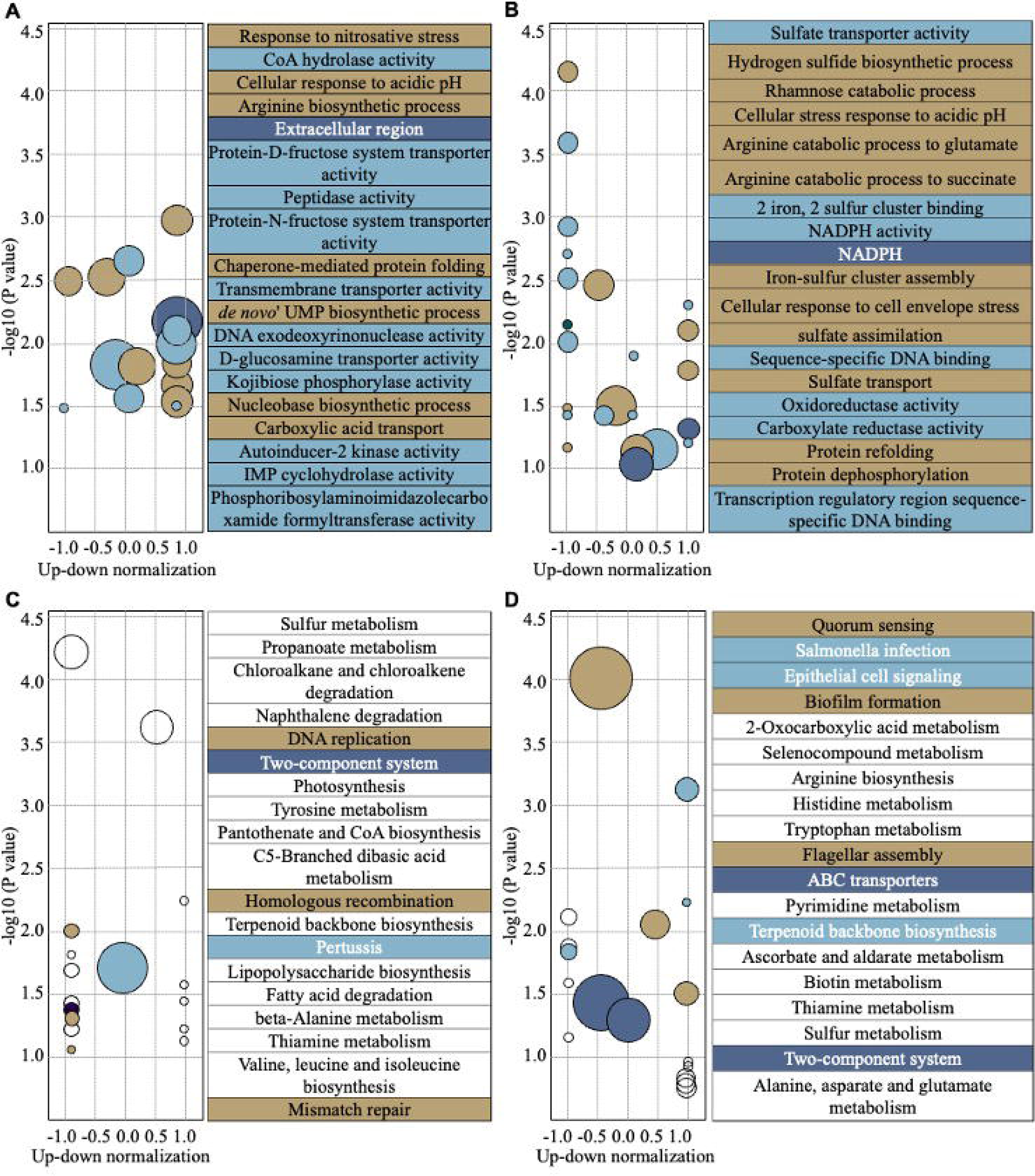
Transcriptional responses of the wild type and Δ*recA* strain after single treatments with ampicillin for 8 hours. GO analysis was performed following the GOseq approach, and different genes in the wild type **(A, left)** and the Δ*recA* strain **(B, left)** were plotted. KEGG pathway enrichment was assigned according to the KEGG database, and different genes in the wild type **(C, left)** and the Δ*recA* strain **(D, left)** were plotted. The top 20 enrichment pathways are listed in the GO and KEGG enrichment analysis **(A-D, right)**. Total RNA-seq was performed with three repeats in each group.

**Supplementary File 1. Table S1. Other mutations detected in the** Δ*recA* resistant isolates

**Supplementary File 2. Table S2. Strains used in this study**

**Supplementary File 3. Table S3. Plasmids used in this study**

**Supplementary File 4. Table S4. Primers used in this study**

