## Supplementary material for "Fast Evolution of SOS-Independent Multi-Drug Resistance in Bacteria": Table S1

**Table S2. Strains used in this study**

| **Strain** | **Relevant Genotype** | **Parent strain** | **Source** |
| --- | --- | --- | --- |
| DH5α | - | - | Lab stock |
| *E. coli* MG1655 | *recA^+^ lexA^+^* | - | Lab stock |
| RW1570 | *recA^+^ lexA3* | MG1655 | Lab stock |
| JW0612-1 | *recA^+^ lexA^+^ ΔcitB* | MG1655 | Keio Collection |
| JW0611-1 | *recA^+^ lexA^+^ ΔcitA* | MG1655 | Keio Collection |
| JW1134-1 | *recA^+^ lexA^+^ Δymfm* | MG1655 | Keio Collection |
| JW0059-1 | *recA^+^ lexA^+^ ΔpolB* | MG1655 | Keio Collection |
| JW0221-1 | *recA^+^ lexA^+^ ΔdinB* | MG1655 | Keio Collection |
| RW120 | *recA^+^ lexA^+^ΔumuDC* | MG1655 | Lab stock |
| EAW13 | *recA^+^ lexA^+^ΔsulA* | MG1655 | Lab stock |
| JW2669-1 | *lexA^+^ ΔrecA* | MG1655 | Keio Collection |
