## Supplementary material for "Fast Evolution of SOS-Independent Multi-Drug Resistance in Bacteria": Table S2

**Table S3. Plasmids used in this study**

| **Plasmid** | **Features** | **Source** |
| --- | --- | --- |
| pJM1071-*recA* | Low copy vector with constitutive *recA* promoter and native *recA* RBS; *recA* cloned in pJM1071 between *NdeI/XbaI* | Lab stock (59) |
| pKD46 | Temperature sensitive replication (*repA101ts*); encodes lambda Red genes | Lab stock |
| pCP20 | Temperature sensitive origin of replication; encodes the FLP recombinase. | Lab stock |
