## Supplementary material for "Fast Evolution of SOS-Independent Multi-Drug Resistance in Bacteria": Table S3

**Table S4. Primers used in this study**

| **Name** | **Sequence** | **Function** |
| --- | --- | --- |
| *recA-FWD* | AAAAAAGCAAAAGGGCCGCAGATGCGACCCTTGTGTATCAAACAAGACGAGAAACGAGAGAGGATGCTCAC | Construction of *recA* deletion mutant |
| *recA-REV* | CAACAGAACATATTGACTATCCGGTATTACCCGGCATGACAGGAGTAAAAGACGTCTAAGAAACCATTATTATCATGAC |  |
