## Supplementary material for "Fast Evolution of SOS-Independent Multi-Drug Resistance in Bacteria": Table S4

**Table S1. Other mutations detected in the *ΔrecA* resistant isolates**

| **Gene** | **Mutation** | **Genomic Position** | **Animo Acid Changes** |
| --- | --- | --- | --- |
| *stfE* | A>AGGTTTTCGAGAGC | 1209618 | p.Val13fs |
| *puuC* | T > G | 1363695 | p.Phe318Cys |
| *cpxA* | T > C | 4104711 | p.Thr89Ala |
